# Antimicrobial resistance in Arctic soils is mediated by competition and facilitation

**DOI:** 10.1101/2023.10.05.561057

**Authors:** Shamik Roy, Robin Dawson, James A. Bradley, Marcela Hernández

## Abstract

Antimicrobial resistance (AMR) is widespread in terrestrial ecosystems. However, the natural processes shaping the spatial and temporal dissemination of AMR in soils are not well understood. We aimed to determine whether, how, and why AMR varies in recently deglaciated pioneer and developing Arctic soils. We showed that antibiotic-resistant genes (ARGs), mobile genetic elements (MGEs), and antibiotic-resistant bacteria (ARB) are abundant, exhibit a non-uniform distribution, and generally increase with soil age. Our analyses suggest a strong positive relationship between soil age and ARG and ARB, which we attribute to increased competition between microbes in older soils. We also observed a weak negative relationship between soil age and ARG diversity mediated by soil organic matter – suggesting facilitation due to the alleviation of nutrient limitation. The microbial processes regulating the spread of AMR in Arctic soils may be further susceptible to the effects of future climate change and human activities.

**Teaser:** The spatial and temporal spread of antimicrobial resistance in Arctic soils is dependent on microbial interactions for nutrients

## Introduction

Soil represents a natural reservoir for antibiotic resistance genes (ARG) that are either intrinsically evolved in bacteria or acquired under selection pressure through mutation, and recombination and/or through horizontal gene transfer (HGT) (*1–3*). ARGs within bacteria primarily act as a deterrent to the antimicrobial products (antibiotics) secreted by competing microbes in anticipation of maximising their nutrient acquisition from the environment, and thereby pre-date anthropogenic antibiotic use (*4*, *5*). However, the use of antimicrobial agents among humans for treating diseases and in livestock production systems has exacerbated natural antimicrobial resistance (AMR) and triggered its subsequent spread across all terrestrial ecosystems (*1*). We are now beginning to understand the molecular and genetic basis of AMR, as well as the spatial boundaries of AMR - with positive detection in remote areas, including the Arctic soils (*2–4*, *6–8*). However, the processes shaping the temporal and spatial dynamics of AMR dissemination in the soil are not well known. In particular, the understanding of whether, how, and why AMR varies during soil development (pedogenesis) remains inadequate, particularly in natural environments with minimal anthropogenic influence (*9*). Addressing this knowledge gap will improve our understanding of bacterial competition, adaptation, and evolution in response to antibiotic pressure, and enable effective management of the current and projected AMR spread through antimicrobial stewardship.

Temporal changes in AMR during pedogenesis can be studied through direct approaches that involve the observation of AMR over time, as well as through indirect approaches that include the use of a space-for-time substitution. Soil formation, for instance, occurs over timescales ranging from decades to millennia. These timescales are often beyond the reach of conventional experiments and direct observation (e.g. via repeated measurements). However, they may be well-suited to observation by indirect means, including a chronosequence approach (*10–12*). The chronosequence approach enables temporal dynamics to be inferred by comparing multiple sites that were formed from the same parent material or substrate and share similar environmental conditions but differ in the time since their formation or at various stages in the soil formation process (*12*). In glacier forefields, soils that are closer to the snout of a retreating glacier are younger, meaning they are at an earlier stage in the soil development process, compared to soils that are distant from the glacier snout and have been exposed for a longer period. A space-for-time substitution can then be employed in the forefields of retreating glaciers to study pedogenesis, including the succession of microorganisms and vegetation (*13*), and other temporally varying factors such as AMR spread (*9*).

Glacier forefields in the Arctic are one of the last remaining frontiers of minimal human activity and influence. However, the Arctic is currently undergoing rapid climate warming, and glaciers are retreating, subsequently exposing pioneer soils (*14*, *15*) and opening new niches for microbes to proliferate - with potential consequences for AMR (*8*, *16*). In Arctic soils, sources of AMR can originate from natural competition among resident microbes, as well as arising through dispersion (for example, by birds, livestock, and humans (*8*). In this context, ARGs, mobile genetic elements (MGEs), and antibiotic-resistant microbes play a role in facilitating the spread of AMR. However, there is a possibility that current and future industrial expansion, as well as an increase in the Arctic population, could also lead to significant quantities of undegraded antibiotics and ARGs entering the natural environment (*17*). Therefore, it is pertinent and timely to investigate whether and how AMR spreads as pioneer soils develop in the Arctic.

In this study, we investigated the trajectory of AMR spread in recently deglaciated Arctic soils. As recently exposed pioneer soils develop, microbes in early-stage (i.e. younger) soils may be taxonomically less diverse and functionally constrained compared to developed (i.e. older) soils (*16*, *18*), as well as subjected to environmental filtering through factors such as nutrient availability (which is generally lower in pioneer soils than in older soils) (*13*, *16*, *19*). The direction and magnitude of inter- and intra-species microbial interactions that regulate nutrient use and the succession of multi-species microbial assemblages, such as competition, facilitation, and complementation, also varies throughout different stages of soil development (*20*). These microbial processes can lead to two contrasting effects (Fig. 1). Firstly, in younger soils, limited nutrient availability can increase competition for nutrients among microbes, which in turn can enhance the prevalence of ARGs and ARB (*21–23*). In contrast, in older soils, the alleviation of nutrient limitation brings complementarity in nutrient use and facilitates interactions between microbes. This, in turn, eases the competition for nutrients, and reduces the abundance of ARGs and ARB. Secondly, in younger soils, environmental filtering of microbes, coupled with limited nutrient accessibility, leads to a lower probability of nutrient use overlap among them. This subsequently reduces competition, resulting in a decrease in the abundance of ARGs and ARB. Conversely, in older soils, taxonomically and functionally rich microbial communities, along with higher nutrient accessibility, exhibit greater nutrient use overlap. This, in turn, leads to narrower realised niches and a decrease in nutrient complementation. It also increases competition through antagonistic interactions among microbes, ultimately resulting in an increase in the abundance of ARGs and ARB (Fig. 1) (*21*, *23*). These conflicting phenomena led us to two opposing expectations where prevalence of ARGs and ARB can either increase or decrease with soil age.

**Figure 1.**
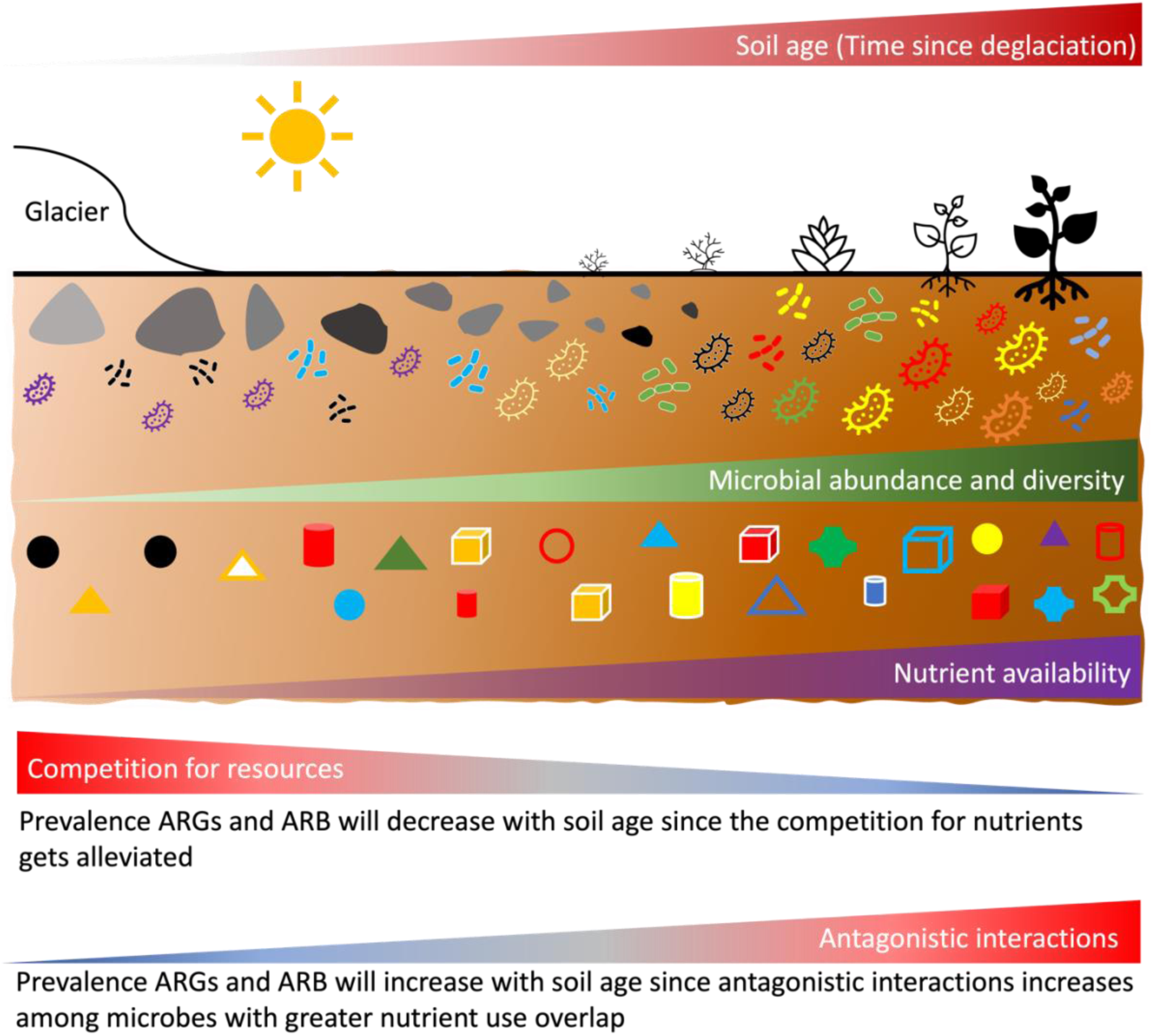
Conceptual illustration of soil development processes following glacier retreat. The triangles indicate the magnitude of variables indicated in the accompanying text, with widening indicating increasing magnitude and thinning indicating decreasing magnitude. The diagram indicates various expectations that, with increasing distance from the glacier snout: age of soil increases, microbial abundance and diversity increase, nutrient availability decreases, competition for nutrients decreases, and antagonistic interactions increase. The complex interplay of processes lead to alternate expectations with regard to the prevalence of ARGs and ARB in different stages of soil development.

Specifically, we tested our competing expectations via three inter-related research questions. First, we assessed whether the forefield of retreating glaciers in the Arctic (Fig. 2) harbours ARGs and ARB, and if so, we investigated their abundance and distribution. Second, we investigated whether the abundance and distribution of ARGs and ARB varies with stages of soil development. Finally, we explored the factors that best explain the distribution of ARGs and ARB in proglacial Arctic soils.

**Figure 2:**
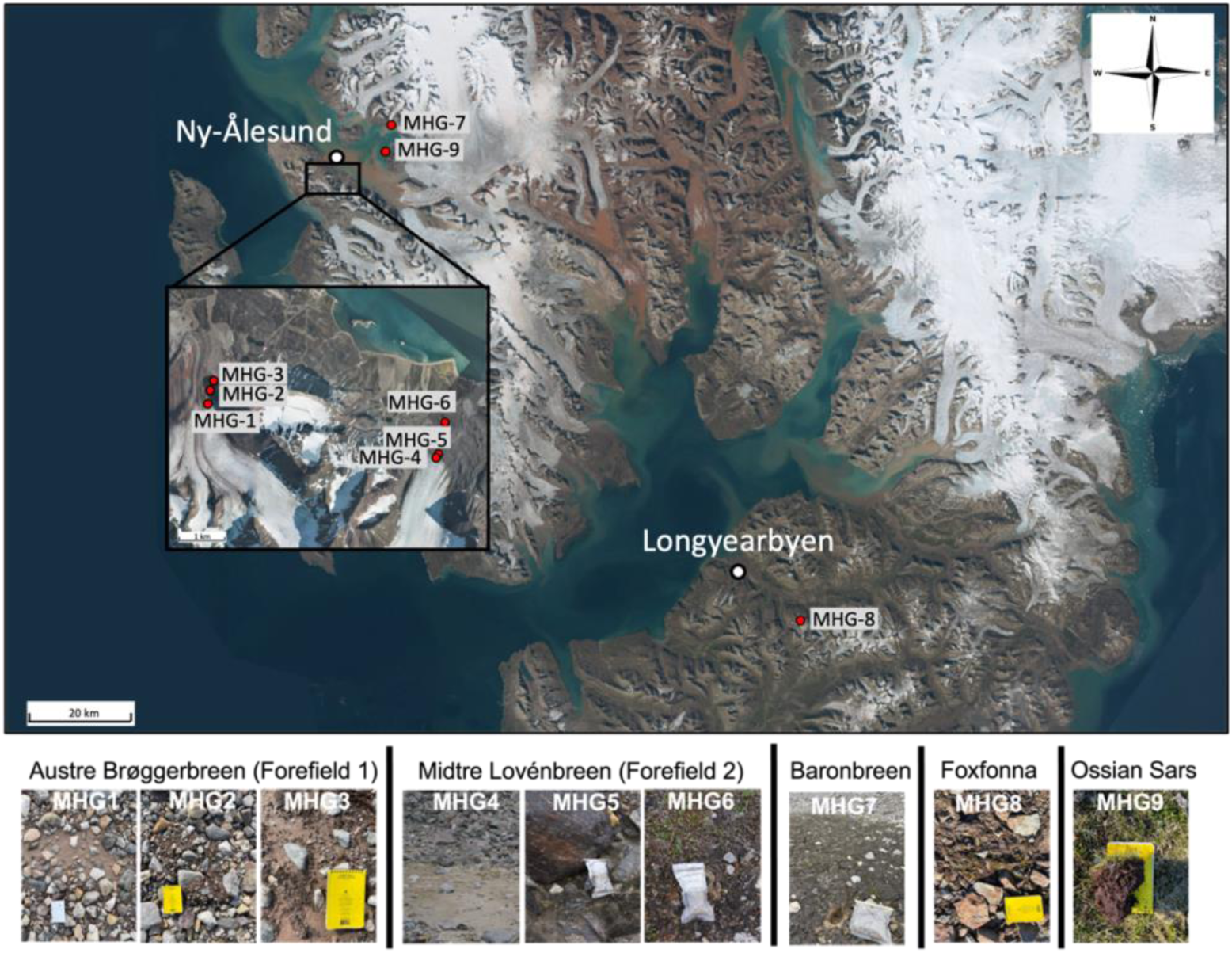
Map of the sampling locations across in Svalbard, and photographs of sampling sites. The white/clear whirlpak bags and field notebooks pictured are approximately 15cm in length.

## Results

### Abundance of antibiotic-resistant genes (ARGs) and mobile genetic elements (MGEs)

qPCR of ARGs belonging to different antibiotic classes showed that not all the ARGs we screened were detected in all soil samples from the glacier forefields. Eight ARGs (*tetA*, *tetX*, *ampC*, *bla_CTX-M_*, *bla_NDM-1_*, *pncA*, *oqxA*, and *vanB*) were present in more than one sample (Fig. 3). No ARGs were detected in pioneer soil (4 years since exposure from glacier retreat) from the forefield of Austre Brøggerbreen (MHG1). Four ARGs were found in developed soils from the Baronbreen forefield (83 yrs, MHG7), and eight ARGs were found in late-stage developed soil approximately 2,000 years old (MHG9). Among the eight positive ARGs, *bla_NDM-1_* was found in four soils, *ampC* was found in six soils, *tetA* and *bla_CTX-M_* were found in seven soils, while the remaining four ARGs were found in eight soils (Fig. 3a). The most dominant ARG was *pncA* (3.17 – 5.40 log_10_ copies g^-1^ soil). This was followed by *vanB* (1.81 – 4.26 log_10_ copies g^-1^ soil) and *tetX* (2.07 – 4.14 log_10_ copies g^-1^ soil). The least abundant ARG was *bla_NDM-1_* (1.74 – 2.76 log_10_ copies g^-1^ soil). We found that the relative abundance of the eight ARGs increased with soil age (p<0.05; Fig. 3 a-b, S1; Table S1). The oldest soil (2,000 years, MHG9) showed the highest relative abundance for all ARGs. However, the rate of increase in ARG abundance with soil age differed among different glacier forefields and overall, across all samples (Fig. 3 a-b, S1, S2; Table S1, S2). We also measured the abundance of two MGEs (*tnpA* and *intI1*) using qPCR. *tnpA* was not detected in any sample (Fig. 3). *intI1* was detected in all soils, including the pioneer soils from the forefield of Austre Brøggerbreen (4 years since exposure, MHG1) in which no ARG was detected. The relative abundance of *intI1* varied between samples and ranged from a minimum of 1.94 log_10_ copies g^-1^ soil in pioneer soil MHG1 (5 yrs), to a maximum of 4.81 log_10_ copies g^-1^ soil in the most developed soil MHG9 (2,000 yrs). Similar to ARG, the relative abundance of *intI1* showed a positive relationship with soil age, indicating that an increase in soil age corresponds to an increase in *intI1* abundance (r=0.74, P=0.001; Fig. 3, S3). The slope of this relationship varied among different glacier forefields and across all samples (Fig. S3).

**Figure 3.**
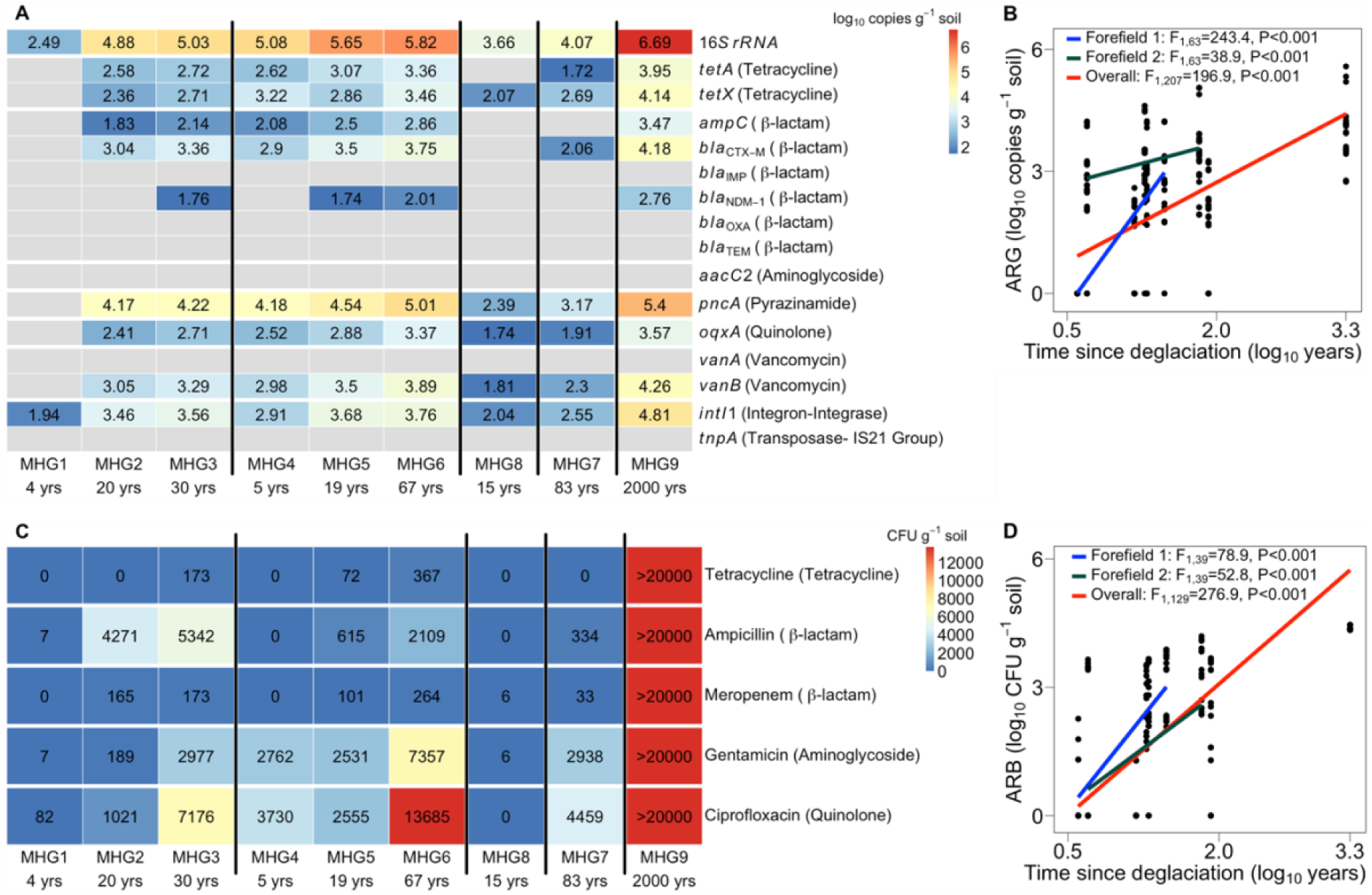
Antimicrobial resistance in deglaciated Arctic soils. (**A**) Abundance of 16S rRNA gene, antibiotic resistant genes (ARGs), and mobile genetic elements (MGEs) in soils sampled from different glacier forefields and stages of development. (**B**) Relationship of ARGs with soil age (time since deglaciation). (**C**) Abundance of antibiotic resistant cultivable heterotrophic bacteria (ARB) in soils sampled from different glacier forefields and stages of development. (**D**) Relationship of ARB with soil age (time since deglaciation). Forefield 1 (blue line) corresponds to Austre Brøggerbreen (MHG1, MHG2, MHG3); Forefield 2 (green line) corresponds to Midtre Lovénbreen (MHG4, MHG5, MHG6); Overall (red line) corresponds to all samples evaluated together. The numbers in each cell denote the absolute abundance of different ARGs (log_10_ copies g^-1^ soil) and ARBs (CFU g^-1^ soil). Grey cells in (A) indicate that ARGs and MGEs were not detected.

### Antibiotic resistant bacteria (ARB)

The abundance of ARB varied for different antibiotics and different soil ages. Overall, ARB abundance was low (in pioneer soils from Austre Brøggerbreen (0-7 CFU g^-1^ soil; 4 years since exposure, MHG1) and Foxfonna (0-6 CFU g^-1^ soil, 15 years since exposure, MHG8) across all antibiotics (Fig. 3c). There were no ARB in developing soils from Midtre Lovénbreen (19 years, MHG5) for all antibiotics except gentamicin. The highest abundance was observed in late-stage developed soils (>9000 CFU g^-1^ soil; 2,000 yrs, MHG9) for all antibiotics. The abundance of ARB for all five antibiotics varied with soil age, such that an increase in soil age corresponded to an increase in ARB abundance (p<0.05; Fig. 3 c-d, S4; Table S3). Similar to ARGs and MGE, the slope of the positive relationship between ARB abundance and soil age varied among different glacier forefields and across all samples (Fig. 3 c-d, S4; Table S3)

Overall, bacteria were most resistant to gentamicin (at a concentration of 16 μg ml^-1^ nutrient medium) in all samples, where the abundance varied from 6 CFU g^-1^ soil in Foxfonna soils (15 yrs, MHG8) to >2,000 CFU g^-1^ soil in six samples. Soils treated with tetracycline (concentration of 16 μg ml^-1^) resulted in no ARB colonies forming in five samples, to >100 CFU g^-1^ soil in three samples (Fig. 3c). For ampicillin (concentration of 32 μg ml^-1^), colony abundance varied from no ARB colonies in two samples, to >2,000 CFU g^-1^ soil in four samples. Similarly, for meropenem (concentration of 4 μg ml^-1^), the abundance of colonies varied from no ARB colonies in two samples to >100 CFU g^-1^ soil in five samples. Finally, for ciprofloxacin (concentration of 1 μg ml^-1^), the abundance of colonies varied from no ARB colonies in two samples to >100 CFU g^-1^ soil in five samples.

### Microbial diversity

Species richness of OTUs ranged between a minimum of 1,152 in pioneer soil from Austre Brøggerbreen (4 yrs, MHG1) to a maximum of 2,930 in developed soils (2,000 yrs, MHG9). Shannon diversity ranged between 4.23 in pioneer soil from Midtre Lovénbreen (5 yrs, MHG4) and 6.14 in developed soil from Baronbreen (83 yrs, MHG7). Species richness showed a positive relation with soil age (P=0.022; Fig. 4a, S5), but Shannon diversity did not show any relationship with soil age (P=0.107; Fig. 4b, S5). The relative abundance of the bacterial phyla varied between samples along the chronosequence with Pseudomonadota being the most dominant phyla across all samples, followed by Actinomycetota and Bacteroidota (Fig. 4c). Among Pseudomonadota, *Shingomnas* was the most identified genus (Fig. 4d).

**Figure 4.**
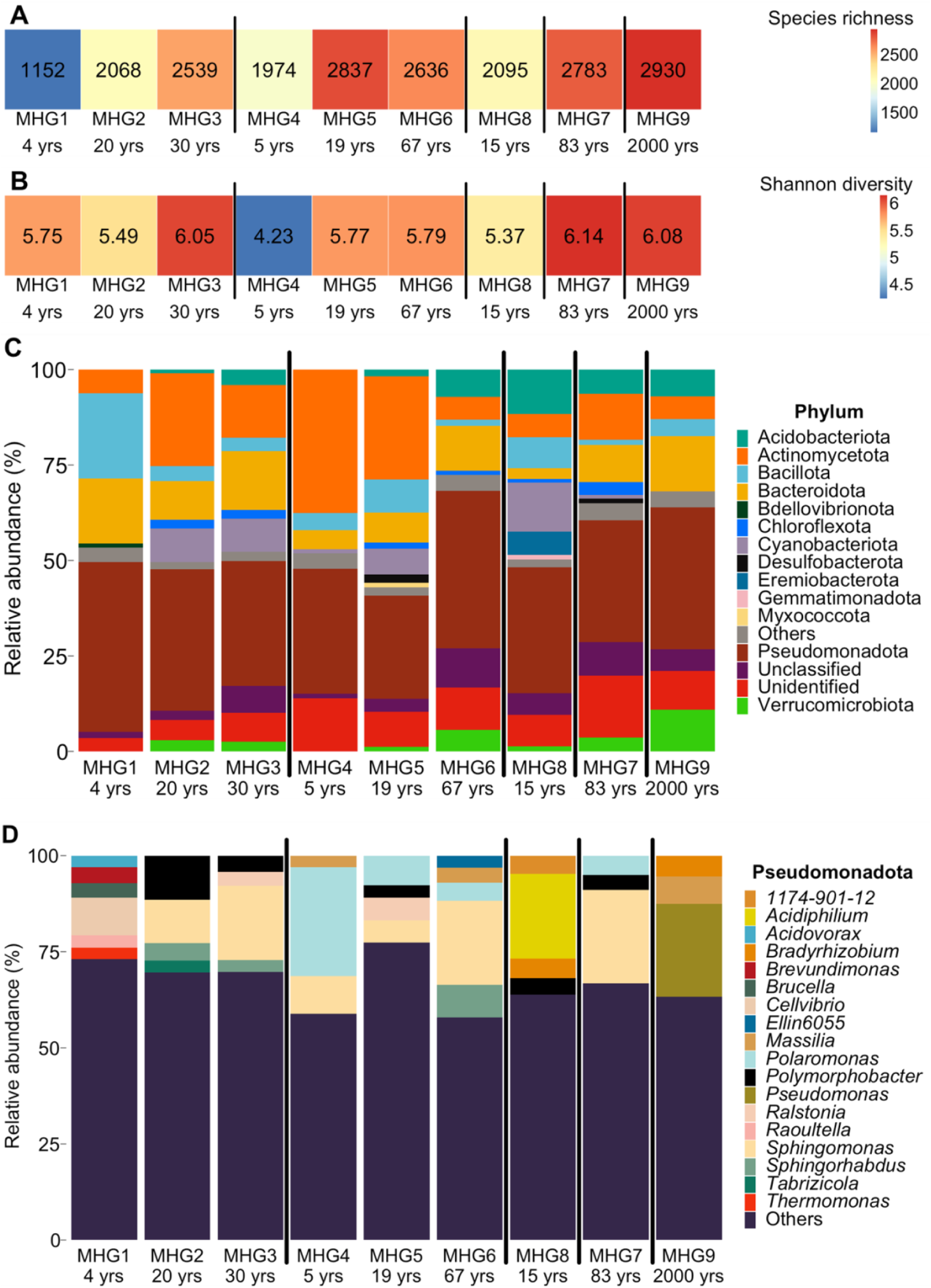
Microbial diversity in Arctic soils from different glacier forefields and stages of development. Microbial diversity is evaluated as (**A**) species richness, (**B**) Shannon diversity, and the relative abundance of dominant (**C**) microbial phyla and (**D**) Pseudomonadota. Numbers in each cell of A and B denote species richness and Shannon diversity, respectively, for each site.

### Inter-relationship between different variables

Partial least squares-path modelling was used to assess the inter-relationship between soil age, edaphic factors, and microbial variables in Arctic soils (Fig. S6, S7; Table S4, S5). The goodness of Fit for the model was 0.692 (Fig. 5), and the model explained 99% of the variation in ARG abundance. Overall, soil age led to an increase in ARG abundance, after accounting for both direct and indirect paths (direct effect (soil age → ARG) was positive with a path coefficient of 0.475 (P<0.001)). There were two main indirect paths: one meditated via diversity and MGE (soil age → bacterial diversity → MGE → ARG) with a net positive effect, and another mediated via edaphic factors (soil age → soil organic matter → ARG and soil age → bulk density → ARG) with a net negative effect. Collectively, the net indirect effect of soil age on ARG was also positive (effect=0.184; Fig. 5). ARG had a positive effect on the abundance of antibiotic-resistant bacterial colonies (ARG → antibiotic resistant bacterial colonies; path coefficient=0.381, *P*<0.05; Fig. 5). Bacterial diversity also had a positive effect on the abundance of antibiotic-resistant colonies (bacterial diversity → antibiotic resistant bacterial colonies; path coefficient=0.321, *P*<0.05; Fig. 5). In this way, soil age had a net positive indirect effect on the abundance of antibiotic-resistant colonies mediated through bacterial diversity and ARG (effect=0.520; Fig. 5).

**Figure 5.**
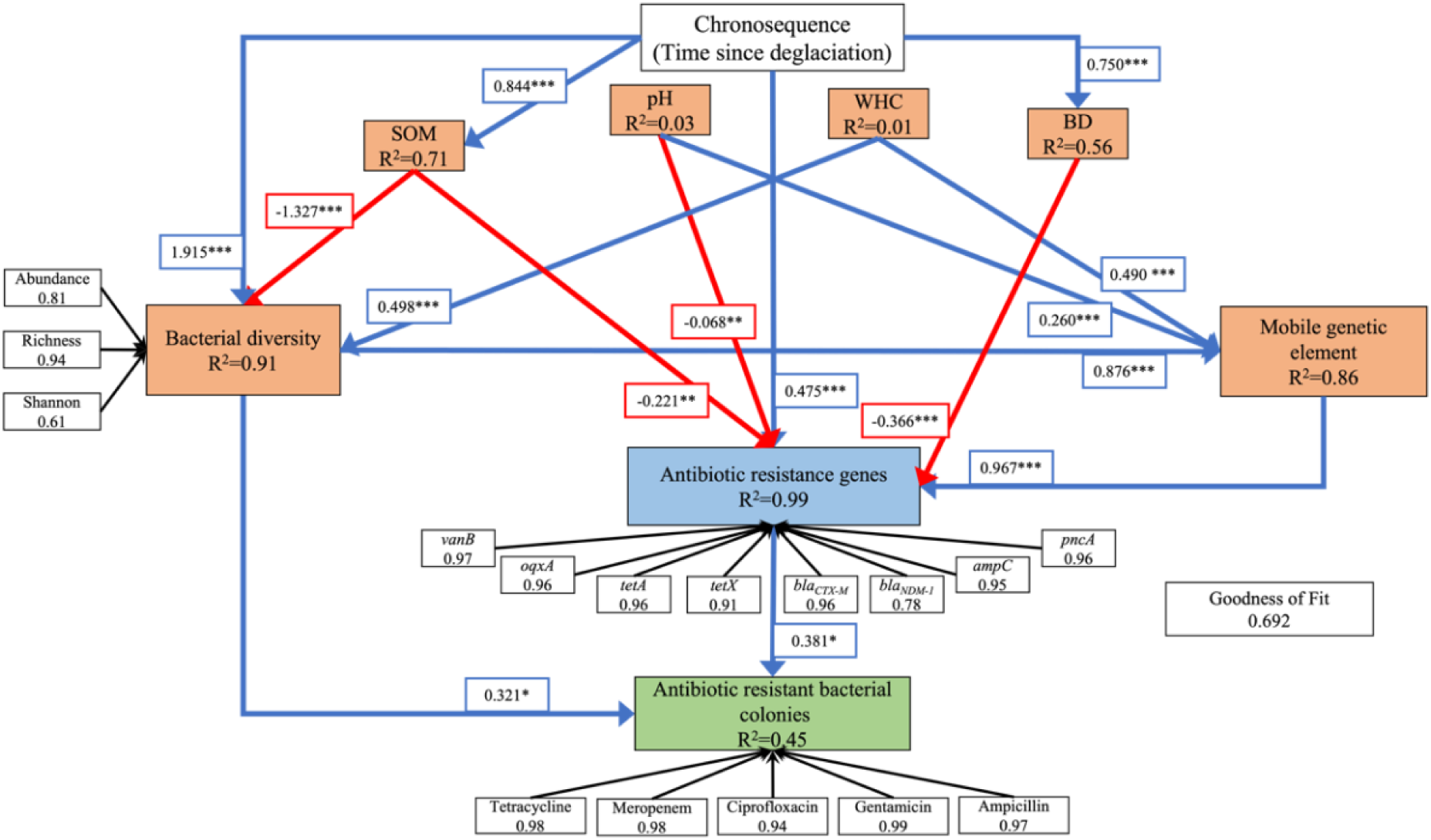
Path analysis using partial least squares path model (PLS-PM) to evaluate inter-relationships among bacterial diversity, mobile genetic element, antibiotic resistant genes, antibiotic resistant bacterial abundance, and soil edaphic factors in developing Arctic soils from deglaciated forefields. The numbers next to the arrows indicate the path coefficients representing the relationships. Path coefficients were calculated after 1,000 bootstraps. The values in the box next to each indicator in the latent variable represent the loadings of the measured variable or indicator. To reduce clutter, only significant paths (α = 0.05) are plotted (for the full model, see companion in Fig. S7 and Table S5). The model is assessed using the Goodness of Fit (GoF) statistic, which is 0.692. SOM: soil organic matter; WHC: water holding capacity; BD: bulk density. Blue arrows represent significant positive relationships, while the red arrows represent significant negative relationships. Black arrows represent the relationship of the latent variable with its block of the indicator. R^2^ values indicate the variance explained by the model for each variable. Asterisks next to each path coefficient represent statistical significance (***P ≤ 0.001, **P ≤ 0.01 and *P ≤ 0.05).

### Antibiotic resistome

Comprehensive antibiotic resistant database (CARD) was used to identify and classify putative ARGs or resistome present in the soil metagenomes that were not quantified by qPCR. Overall, the number of ARGs identified varied between samples, with the highest of 225 ARGs in developing soil from Austre Brøggerbreen (30 yrs, MHG3) and the lowest of 97 in late-stage developed soils (2,000 yrs, MHG9) (Fig. 6a). All the Strict Hits of ARGs were found in six samples (MHG3-4, MHG5-8, Fig. S8). There was only one Perfect hit each in pioneer and developing soil samples from Austre Broggerbreen (MHG1 and MHG2), and developing soils from Midtre Lovénbreen (MHG5) that belonged to the TEM beta-lactamase gene family (Fig. S8). The ARGs identified varied among samples when classified into different categories for the mechanism of resistance (Fig. 6c), drug class (Fig. 6d), and gene family (Fig. 6e). Specifically, the dominant mechanism of resistance across all samples was “antibiotic target alteration” with >80% of the ARGs identified. Similarly, the dominant ARGs detected across all samples were *vanW* and *vanY*, which provide resistance to the glycopeptide drug class. 19 gene families were identified across all soil resistomes. We found a weak negative effect (α=0.10) of soil age on the number of ARGs identified using metagenomics (Fig. 6b).

**Figure 6.**
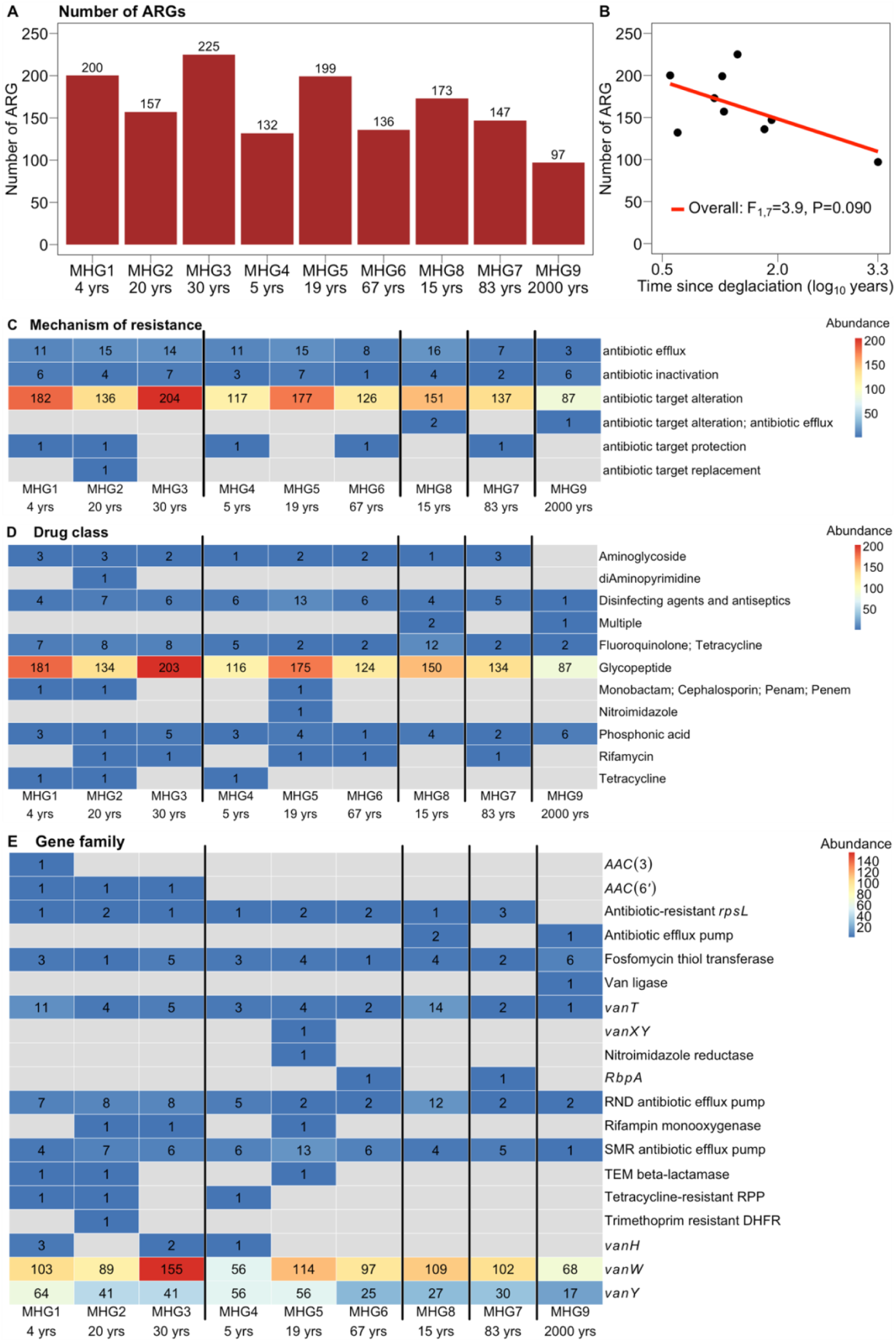
Antibiotic resistome following metagenomics from developing Arctic soils from deglaciated forefields. (**A**) The number of antibiotic resistant genes (ARGs) detected and (**B**) their relationship with soil age (time since deglaciation). (**C**) ARGs identified across categories based on different mechanisms of resistance, (**D**) different drug classes, and (**E**) gene families. The number in each cell denotes the number of ARGs for each soil across different categories. Grey cells indicate the absence of detected ARGs. rbp: RNA-polymerase binding proteins, RND: Resistance-nodulation-cell division, SMR: Small multidrug resistance, RPP: Ribosomal protection proteins, DHFR: Di-hydro folate reductase.

## Discussion

The processes shaping the temporal and spatial dynamics of AMR dissemination in soils are not understood. Here, our aim was to investigate how and why AMR varies in pioneer and developing Arctic soils with minimal anthropogenic influence, using a space-for-time substitution approach. We assessed the prevalence of ARGs and ARB at different stages of soil development in glacier forefields. We found that forefields of retreating glaciers in the Arctic harbour ARGs and ARB, which is consistent with other studies in the Arctic and Antarctic glaciers (Fig. 3) (*8*, *24*, *25*). However, only eight out of the 13 ARGs assessed were present across samples in the chronosequence, and they, along with ARB, exhibited a non-uniform distribution (Fig. 3). The abundance of the remaining five ARGs could be below the detection limit of our protocol or absent from the soils analysed. Furthermore, their abundance increased with soil development stages, from newly deglaciated pioneer soils to more developed soils (Fig. 3, S1; Table S1). Our PLS-PM analyses showed that soil age had both direct and indirect effects on the ARGs (Fig. 5). Finally, we also found a positive relationship between *intI1* abundance, a mobile genetic element facilitating HGT, and all eight of these ARGs (Fig. 5, S6; Table S4). HGT is known to be a major driver of AMR spread among diverse microbial species in the community (*1*, *3*). These findings may have implications for how we manage future medical and environmental impacts of AMR spread through time (*26*).

The results from this study strongly support the hypothesis that as soil age increases, antagonistic interactions among microbes competing for the same resource leads to an increased prevalence of ARGs and ARB (Fig. 3, 5, S1, S4; Table S1, S3). We propose that as soils undergo pedogenesis, there are corresponding increases in microbial diversity, nutrient use overlap, and competition between microbes to acquire nutrients from their environment (Fig. 1). The antagonistic competitive interaction between soil microbes gives rise to the production of chemically diverse antimicrobial products in the soil environment, further propelling the evolution and spread of genetic countermeasures in the form of ARGs and MGE that provide protection to the microbes against these antimicrobial products. Therefore, as soils develop, the abundance of ARGs and ARB increases. Our alternative expectation was that ARG and ARB abundance would decrease with soil age, due to the alleviation of both nutrient limitation and competition to acquire nutrients between microbes during pedogenesis. Alleviation of nutrient limitation can increase complementarity for nutrient use and reduce nutrient use overlap, which leads to facilitation rather than competition among microbes. Consequently, the necessity for antimicrobial products and corresponding ARGs to provide protection is reduced in mature soils compared to younger soils. Interestingly, our results also support this alternate expectation (Fig. 5), since we found that soil age can negatively affect ARGs and ARB via soil organic matter (see Fig 5). Although this result contradicts the dominant positive relationship observed between ARGs and ARB with soil age, our finding is consistent with a global meta-analyses of soil chronosequences across different ecosystems and environments which found that soil age can have both negative and positive effects on ARG abundance (*9*). This apparent contradiction has been attributed to the relative strength of competition and facilitation between microbes during soil development. Therefore, we posit that over the course of pedogenesis in Arctic soils, even though competition and facilitation among microbes in natural communities are working in tandem, showing both positive and negative effects of soil age on ARGs and ARB, the effect of competition is dominant.

Microbial interactions (competition and facilitation) are important factors in shaping the temporal variation of ARGs and ARB (*1*, *23*). The proximate determinants for the development of antibiotic resistance and its subsequent spread are horizontal gene transfer and mutation, as well as diverse microbial assemblages that can harbour ARGs (*3*, *5*). In fact, we showed there is variation in ARGs and ARB with soil age in the Arctic glacier forefields, and this variation can be meditated through microbial diversity and HGT (Fig. 5, S6; Table S4). Furthermore, we showed that these proximate determinants are not mutually exclusive (Fig 5), since diverse microbial assemblages can support a higher MGE abundance and, accordingly, greater rates of HGT. These results provide strong empirical support to the processes that were previously hypothesised for temporal AMR dissemination in soil (*1*, *5*, *23*).

Our results also highlight that the rates and trajectories of AMR spread in the Arctic are spatially heterogeneous. Differences in the abundance of ARGs and ARB with soil age between the two glacier forefields with similar soil age profiles (Fig. 3, S1–S5; Table S1–S3) may arise from varying initial levels of AMR (including ARG abundance and microbial diversity) as well as inherent variability in soil characteristics between different glacier forefields (*16*, *27*, *28*). It has been hypothesised that the initial microbial diversity in recently deglaciated soils results from spatial variation in the paleoenvironment before and during glacier formation (*29*). We found clear differences in pioneer soils between Midtre Lovénbreen – harboring seven out of the 13 examined ARGs and higher richness (1974 OTUs), and Austre Brøggerbreen - which does not harbour any of the examined ARG and had a lower richness (1152 OTUs). It is also well-established that soil edaphic variables can determine the relative importance of competition and facilitation in microbial assemblages, thereby affecting AMR. Edaphic variables varied between the different glaciers as well as between the stages of soil development – which directly and indirectly influenced the abundance of ARGs and ARB (Fig. 5, Table S6). These soil edaphic variables can influence both the proximate and ultimate determinants of AMR through habitat filtering (*3*, *9*, *30*).

Our metagenomic analyses of developing Arctic soils revealed a prevalence of *vanW* and *vanY* gene families, genes that are attributed to antimicrobial resistance by altering the target of glycopeptide drug class antibiotic Vancomycin (Fig. 6). Along with the positive identification of *vanB*, the prevalence of *vanW* and *vanY* in pioneer Arctic soils that are largely devoid from human influence suggests that vancomycin resistance in the soil might be ancient and predate clinical antibiotic usage. This interpretation is consistent with a study that discovered the presence of functional vancomycin-resistant genes in the ancient DNA extracted from 30,000-year-old Beringian permafrost sediments (*4*).

We found a negative effect of soil age on the number of ARGs (Fig. 6), which we suspect may result from a combination of two factors. First, over the course of soil development, the diversity of ARGs may decrease due to environmental filtering and reduced competition among microbes for resources, with only selected ARGs remaining prevalent in older soils. Secondly, the apparent negative effect of soil age on the number of ARGs may be an artefact of methodological limitations: as microbial diversity increases, there is a reduction in read coverage and sometimes an increase in non-uniform read coverage in metagenomic data (*31*). The highly diverse and large metagenomes of developed soils (compared to pioneer soils) may introduce noise from viral DNA and eukaryotic DNA, which can dilute the reads from bacteria and lead to an uneven representation or reduced prevalence of certain reads or contigs during the bioinformatic analyses.

Our study may also be subject to other methodological and practical limitations. First, we did not assess the contribution of mutations towards the abundance and diversity of ARGs. Second, we did not account for the quality and bioavailability of soil nutrients, which we suspect may influence AMR spread through changes in microbial interactions and diversity (*32*). Third, we did not consider other soil properties, such as redox status and soil moisture. Additionally, the outcomes of the path model do not imply causation; instead, they point toward testable hypotheses and raise new questions to identify the determinants of AMR and its spread along the chronosequence.

In summary, we elucidated the temporal patterns of AMR spread in newly developing Arctic soils. We revealed that ARGs, MGE, and ARB are abundant, have a non-uniform distribution, and generally increase with soil age in Arctic glacier forefields. We conclude that this temporal pattern of AMR is a consequence of the intricate and dynamic balance between microbial competition and facilitation. Retreating Arctic glaciers are exposing pioneer soils that undergo pedogenesis and are also subject to the effects of climate change and human activity. The fundamental microbial processes that regulate the spread of AMR may be further susceptible to the effects of future climate change and human activities in the Arctic.

## Material And Methods

### Study area and sampling

We sampled soils in the forefields of four high-Arctic glaciers in Svalbard (Fig. 2): Austre Brøggerbreen, Midtre Lovénbreen, Baronbreen, and Foxfonna. Soils were sampled from three locations at different distances from the glacier snout in two glacier forefields: Austre Brøggerbreen (sampled in June 2022: MHG1, 2 and 3, with soil ages of 4, 20 and 30 years respectively), and Midtre Lovénbreen (sampled in July 2021: MHG4, 5 and 6, with soil ages of 5, 19 and 67 years respectively). In Baronbreen (July 2021: MHG7, 83 years) and Foxfonna forefields (August 2021: MHG8, 15 years), soils were sampled from one location each. Soil was also sampled (July 2021)from a location representing late-stage developed soils (MHG9, 2,000 years). Samples were stored frozen at -20^°^C. Soil age (i.e. time since deglaciation) was approximated using aerial photography and satellite imagery available from TopoSvalbard (NPI/USGS Landsat) and Sentinel Hub (https://www.sentinel-hub.com, Sinergise Ltd), as well as data from Bourriquen *et al*. (2018) (*33*).

### Edaphic factors

Soil organic matter (SOM), pH, water holding capacity, and bulk density were measured following standard procedures (*34*) (Table S6, *see additional methods in the SI for more details*).

### Antibiotic resistant bacteria (ARB)

The enumeration of culturable ARB in soil samples was performed using the spread plate method. Briefly, soil samples were thawed at 4°C for 5 hours, and suspended in sterile water at 1:2 (w/v) ratio to prepare slurry that acted as an inoculum. 0.1 ml of the soil slurry was spread in triplicates onto tryptone soy agar plates supplemented with cycloheximide (60 μg ml^-1^) alone as a control to prevent fungal growth, or cycloheximide (60 μg ml^-1^) plus one of the following antibiotics: tetracycline (16 μg ml^-1^; tetracycline antibiotic class), ampicillin (32 μg ml^-1^; β-lactam class), meropenem (4 μg ml^-1^; β-lactam class), gentamicin (16 μg ml^-1^; aminoglycoside class), and ciprofloxacin (1 μg ml^-1^; quinolone class). These antibiotics encompass different antibiotic classes (Table S7) and are frequently used in human and veterinary medicine against a range of pathogens as different lines of defence. The concentration of each antibiotic was selected in accordance with the Clinical & Laboratory Standards Institute (CLSI) and the European Committee on Antimicrobial Susceptibility Testing (EUCAST) guidelines that recommend the antibiotic concentration in which different pathogens become resistant. The plates were then incubated for 10 days at 4°C. Cycloheximide was added to all the plates to prevent fungal growth. After incubation, the number of culturable ARB was calculated as colony-forming units (CFU) for both control and antibiotic-supplemented plates and represented as CFU g^-1^ soil.

### Soil DNA extraction

DNA was extracted from soil following a protocol that includes the CTAB method followed by a cleaning step using Qiagen DNeasy PowerSoil Pro Kit (*35*). Extracted DNA was stored at -20°C for further molecular studies. These included: 1) qPCR to determine the abundance of selected ARGs and MGEs, 2) 16S rRNA gene sequencing for community composition, and 3) metagenomic sequencing to characterise the antibiotic resistome of soil samples along the chronosequence (*see additional methods in the SI for more details*).

### qPCR of 16S rRNA gene, ARGs, and MGE

The abundance of the 16S rRNA gene, 13 different ARGs (Table S7, S8) and two MGE genes were quantified by real-time qPCR using a StepOne Plus real-time PCR instrument (Applied Biosystems). The ARGs represented resistance to tetracycline (*tetA, tetX*), ampicillin (*ampC*), beta-lactams (*bla_CTX-M_, bla_IMP_, bla_NDM-1_, bla_OXA_, bla_TEM_*), vancomycin (*vanA, vanB*), quinolone (*oaxA*), pyrazinamide (*pncA*), and aminoglycoside (*aacC2*). The two MGEs were integron (*intI1*) and transposase (*tnpA*), which are major determinants of HGT in Arctic soils (*8*). The qPCR assay was performed in triplicate for each target gene, along with a control (no DNA). For quality control measures, a sample is positive for ARG and MGE if the threshold cycle (Ct) is < 29, and the Ct value was < 29 for at least two of the three replicates. Quantification of 16S rRNA gene abundance was done by generating a standard curve through serial dilutions (10^-1^ to 10^-5^) of the genomic DNA of *Variovorax* sp. WS11 (*36*). 16S rRNA gene abundance was calculated against their respective standard curve and expressed as copies per gram of soil. ARG and MGE abundance were quantified by calculating relative gene copy numbers using the following equation: *Gene copy number* = 10^(29−*Ct*)⁄(10⁄3)^, where Ct is the threshold cycle for each PCR reaction and 29 is the limit of the threshold cycle for a positive reaction (*37*). The abundance of ARG and MGE are expressed as copies per gram of soil (*see additional methods in the SI for more details*).

### Community profile

Microbial Operational Taxonomic Units (OTU) were estimated following amplification and sequencing of the 16S rRNA hyper variable region V3-V4 for all the DNA samples.

Microbial diversity was expressed as species richness and Shannon diversity (*see additional methods in the SI for more details*).

### Metagenomics

ARGs were also identified from the metagenome of individual soil samples. Briefly, the extracted DNA was sequenced on an Illumina NovaSeq 6000 using a paired-end dual-indexed run. The reads from metagenome sequencing were assembled into longer contiguous sequences (scaffolds). The assembled scaffolds were then screened for ARGs against the comprehensive antibiotic resistance database (CARD) using the resistance gene identifier (RGI) software version 6.0 (*38*). RGI performs predictions of complete open reading frames (ORFs) using Prodigal 2.6.3 (*39*), followed by alignment of these ORFs with CARD reference proteins using DIAMOND 0.8.36 (*40*) (*see additional methods in the SI for more details*).

### Data analyses

To analyse the temporal variation in ARGs and ARBs, linear mixed-effect statistical modelling was performed. Soil age (i.e. time since deglaciation, years; continuous variable) was used as an explanatory variable (i.e., fixed-effect). The identity of each ARG and antibiotic (for ARB) was modelled as the random-effect. This was done separately for samples from Austre Brøggerbreen (MIG1, MHG2, MHG3), Midtre Lovénbreen (MHG4, MHG5, MHG6), and for all samples together (overall). Subsequently, the temporal variation in individual ARGs, ARB for each antibiotic, MGE, and diversity parameters were analysed using linear models with soil age or time since deglaciation (in years; continuous variable) as the predictor variable. A key assumption of linear mixed-effect models is that the residuals are normally distributed. This assumption was verified using observed and theoretical quantiles of residuals for all variables (qq-plot). Data on ARGs, ARB, MGE, and soil age required a log_10_-transformation to meet this assumption. The analyses were performed in R 4.1.1 (*41*) using nlme library (*42*).

The inter-relationships between different microbial variables along the chronosequence were assessed with partial least squares path modelling (PLS-PM). PLS-PM helps evaluate paths with complex multivariate relationships among observed and latent variables (*43*). PLS-PM is advantageous for our study because it does not require any normality assumptions for the data, which is a pre-requisite for variance-covariance based structural equation model (SEM). Furthermore, PLS-PM performs well in the analyses of complex models using smaller samples. Additionally, PLS-PM is an exploratory approach that helps generate new hypotheses on the processes involved as opposed to the SEM, which tests the validity of the a-priori hypothesised pathways. The analyses were done in R 4.1.1 (*41*) with plspm library (*44*) (*see additional methods in the SI for more details*).

## Acknowledgements

We thank Andrew Baughan from high-performance computing cluster (ADA) at UEA for his assistance with metagenomics analyses. We thank the AMBER-ICE, SUN SPEARS, and WAVES field teams, as well as the Ny-Ålesund Research Station – Sverdrup, the UK Arctic Research Station, and the Czech Arctic Research Station in Svalbard for assistance in the field.

## Funding

Natural Environment Research Council Discipline Hopping (DH) for Discovery Science grant NE/X018180/1 (SR, MH)

Royal Society Research Fellows Enhanced grant RF\ERE\210050 (MH)

Royal Society Dorothy Hodgkin Research Fellowship DHF\R1\211076 (MH)

Natural Environment Research Council (NERC) grant NE/T010967/1, NE/V012991/1 (JAB)

NERC COVID Recovery Support Fund (JAB)

The research leading to these results has received Trans-National Access from the European Union’s Horizon 2020 project INTERACT, under grant agreement No. 730938

Sequencing was supported by an award from the Natural Environmental Research Council (NERC) Environmental Omics Facility (NEOF) to MH, JAB

## Author contributions

Conceptualization: SR, MH

Formal analysis: SR, MH

Investigation: SR, JAB, RAD

Methodology: SR, MH, JAB

Resources: MH, JAB

Visualization: SR

Supervision: MH

Project administration: MH

Funding acquisition: MH, JAB

Writing—original draft: SR, MH

Writing—review & editing: SR, RAD, JAB, MH

## Competing interests

All other authors declare they have no competing interests

## Data availability

Sequence data were deposited in the NCBI Sequence Read Archive (SRA) under the bioproject accession numbers PRJNA1019501 for metagenomic raw data and PRJNA1019681 for 16S rRNA gene sequencing. AMR data will be available on request.

## Notes

### Competing Interest Statement

The authors have declared no competing interest.

