## Supplementary materials for "Antimicrobial resistance in Arctic soils is mediated by competition and facilitation"

This supplement supplementary information includes:

Additional methods

Figures. S1 to S8

Tables S1 to S8

Additional References

### Additional Methods

*Edaphic factors*

Soil organic matter (SOM) was determined by measuring the weight loss on ignition for 4 g of soil heated to 450 °C for 4 h. The SOM content was calculated as the difference between the initial and final weights. Soil pH was measured from 1:2.5 (w/v) soil in water suspension using a table-top pH meter. Bulk density was determined by measuring the volume occupied by 4 g soil. Water holding capacity was assessed gravimetrically using a 1:5 (w/v) soil-water suspension.

*Soil DNA extraction*

1 g of soil was mixed with 5 ml of lysis buffer pre-heated at 65 °C and 2.5 µl of 20 mg/ml Proteinase K (*1*). The lysis buffer was prepared by mixing 7.1 g $Na_2HPO_4$, 43.8 g NaCl, 5 g CTAB, 50 ml 1 M Tris (pH 8), 100 ml 0.5 M EDTA and nanopure water to complete 500 ml. Soil slurry in lysis buffer was thoroughly mixed through the vortex and incubated for 2 hours at 65 °C with brief shaking every 15 mins to resuspend the soil. Following incubation, 2 µl of RNase A (10 mg/ml) was added to the mixture and incubated again for 45 mins at room temperature. Then, the mixture was centrifuged at 4000 x g for 10 minutes. Next, the supernatant was transferred to a fresh tube and centrifuged again at 11,000 x g for 20 minutes. The clean supernatant with minimal soil particles was transferred to a new tube, and one volume of 100% ethanol was added to the supernatant. The mixture was sequentially loaded, 700 µl at a time, onto the microfuge silica column (also known as a spin column) from the Qiagen DNeasy PowerSoil Pro Kit. The column was then centrifuged at 15,000 x g for 1 min, and the flow-through was discharged. This step was done until the entire mixture was passed through the column. The column was then treated according to the following up steps recommended by the manufacturer until DNA elution. The eluted DNA was very brown at this point. This eluted DNA will be unsuitable for PCR due to inhibition by humic acid. Therefore, the eluted DNA was washed three times following the manufacturer's protocol in the DNeasy PowerClean Pro Cleanup Kit (Qiagen) to clean the DNA further.  The DNA solution was then loaded onto 0.8% agarose gel to check the integrity. DNA purity was assessed using NanoDrop® Spectrophotometer ND-1000 (Thermo Fisher Scientific, USA), and DNA was quantified following protocol for Qubit high-sensitivity dsDNA kit. Overall DNA yield was in the range of 71-1555 ng $g^{-1}$ soil, with the lowest for MHG1 and highest for MHG9.

*qPCR of 16S rRNA gene, ARGs, and MGE*

The 10 µl qPCR reaction mixture contained 5 µl of 2× SensiFAST™ SYBR® No-ROX qPCR master mix (Meridian Bioscience), 0.6 µl of each primer (10 µM), 2.8 µl sterilised nuclease-free double distilled water, and 1 µl of soil DNA template. The primers used for all the genes are described in Table S8. The protocols of qPCR for different genes were different. For the 16S rRNA gene, amplification was initiated by denaturation at 95 °C for 3 min followed by 40 cycles of denaturation at 94 °C for 30 s, annealing at 50 °C for 30 s, and extension along with signal acquisition at 68 °C for 30 s. For *oqxA* and *tetX* genes, amplification was initiated by denaturation at 95 °C for 3 min followed by 40 cycles of denaturation at 95 °C for 30 s, annealing at 58 °C for 30 s, extension at 72 °C for 30 s, and signal acquisition at 83 °C for 10 s to generate the melting curve. For the rest of the genes, amplification was initiated by denaturation at 95 °C for 3 min followed by 40 cycles of denaturation at 95 °C for 30 s, annealing at 60 °C for 30 s, extension at 72 °C for 30 s, and signal acquisition at 83 °C for 10 s.

*Community profile*

DNA extracted from soil was amplified with primers 341F (5'-CCTAVGGGRBCCASCAG-3') and 806R (5'-GGACTACNNGGGTATCTAAT-3'). Sequencing was done using Illumina PE250 at Novogene, UK. High-quality amplicon sequences were analysed using Qiime1 (*2*). Briefly, for every sample, chimeras were removed using reference-based chimera checking following VSEARCH 2.16.0 (*3*). The contigs were then clustered into OTUs using UPARSE (*4*). Taxonomy was assigned using SILVA_v138 database (*5*).

*Metagenomics*

The metagenomic analysis involved two steps: quality check and assembly. Initially, the quality of the sequenced metagenomic DNA reads for each sample were assessed using FastQC version 0.11.8 (*6*). Any low-quality reads (length <250 bp) were excluded from the analysis using BBDuk version38.68 (*7*). This process resulted in clean, high-quality reads (>250 bp), which were then used for assembly. The processed high-quality reads were then assembled de novo into longer contiguous sequences (scaffolds) using the metaSPAdes version 3.13.1 assembler (*8*). Subsequently, the annotation of ORFs into ARGs using CARD was carried out under the Perfect and Strict paradigms of RGI. The alignment was 'Perfect' when the ORFs matched completely (100%) with the curated reference sequences in CARD. The alignment was 'Strict' when the ORFs fell within the curated BLAST bit score cut-offs, allowing for variation in ORFs from the CARD reference sequence. This is particularly useful for detecting previously unknown variants of known AMR genes or altered antibiotic targets. For this study, we included both Perfect and Strict annotations. The analyses were done on a high-performance computing cluster (ADA) supported by the Research and Specialist Computing Support Service at the University of East Anglia (Norwich, UK).

*Data analyses*

The path diagram for the model that tests the relationships between microbial variables is shown in Figure S7 and Table S5. The model was bootstrapped 999 times to estimate the precision of the PLS parameter estimates. Path coefficients and their statistical significance ($\alpha=0.05$) were reported. Measures of unidimensionality, Cronbach's alpha and DG-rho were also confirmed to be >0.7 for all the latent variables, which indicates that the block of indicators was performing well to measure their corresponding latent constructs. Loadings of each indicator in all the latent variables were >0.7, except Shannon diversity, which indicates that the variability in the indicators was well captured by its latent construct. The quality of the path model was assessed by $R^2$ values for endogenous latent variables and the goodness of fit index that examines the overall prediction performance of the model.

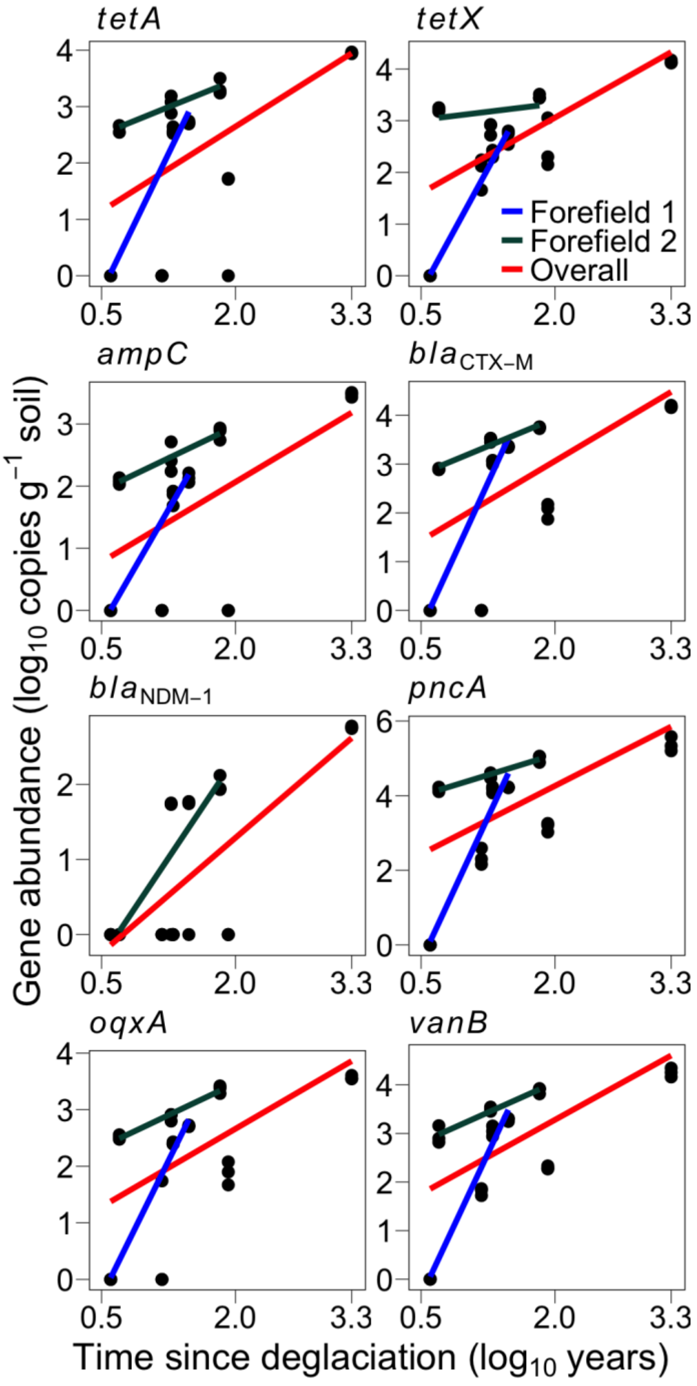


Figure S1. Relationship assessed as regression between chronosequence (time since deglaciation), represented as a continuous variable, and abundance of eight ARGs expressed as copies $g^{-1}$ soil for two glacier forefields and overall, all samples together (red line). The associated statistics are described in Table S1. If the relationship is not significant for any of the combinations, then the regression line (coloured lines) is absent. Forefield 1 (blue line): Austre Brøggerbreen glacier forefield contains samples MHG1, MHG2, MHG3; Forefield 2 (green line): Midtre Lovénbreen glacier forefield contains samples MHG4, MHG5, MHG6.

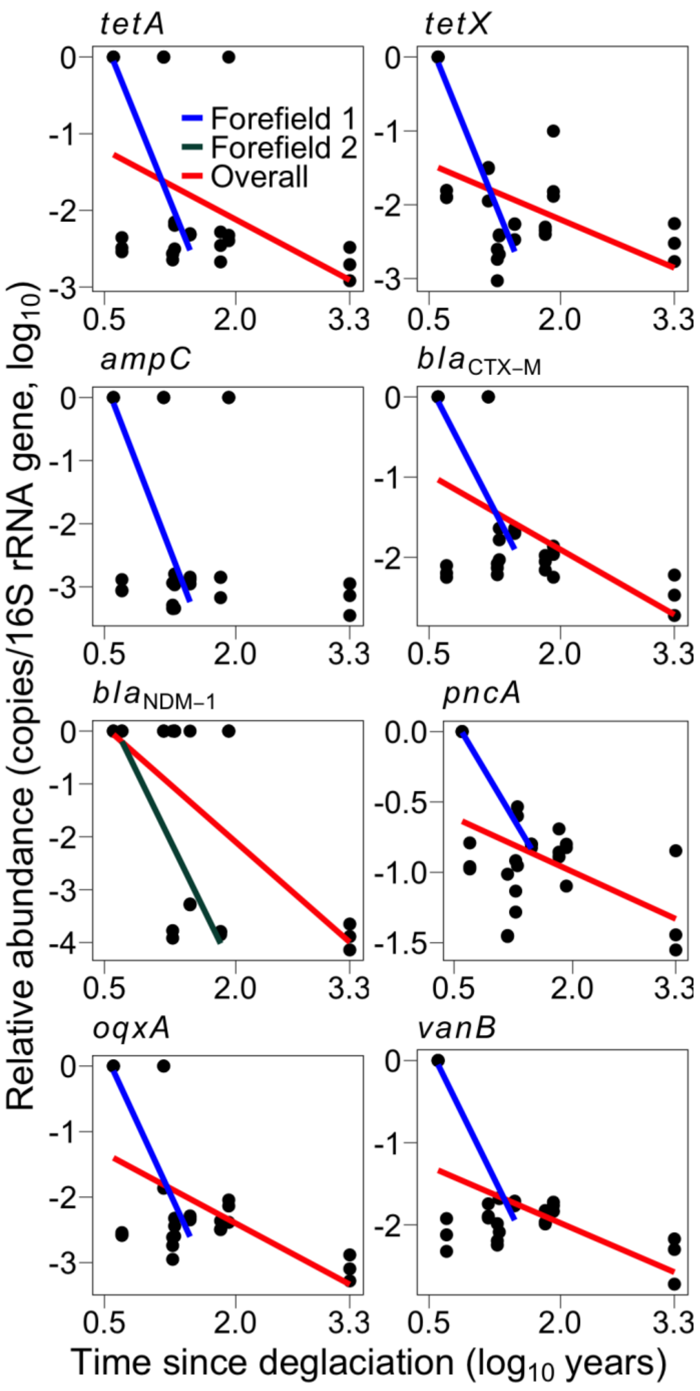


Figure S2. Relationship assessed as regression between chronosequence (time since deglaciation), represented as a continuous variable, and relative abundance of eight ARGs expressed as copies per copy of 16S rRNA gene for two glacier forefields and overall all samples together (red line). The associated statistics are described in Table S2. If the relationship is not significant for any of the combinations, then the regression line (coloured lines) is absent. Forefield 1 (blue line): Austre Brøggerbreen glacier forefield contains samples MHG1, MHG2, MHG3; Forefield 2 (green line): Midtre Lovénbreen glacier forefield contains samples MHG4, MHG5, MHG6.

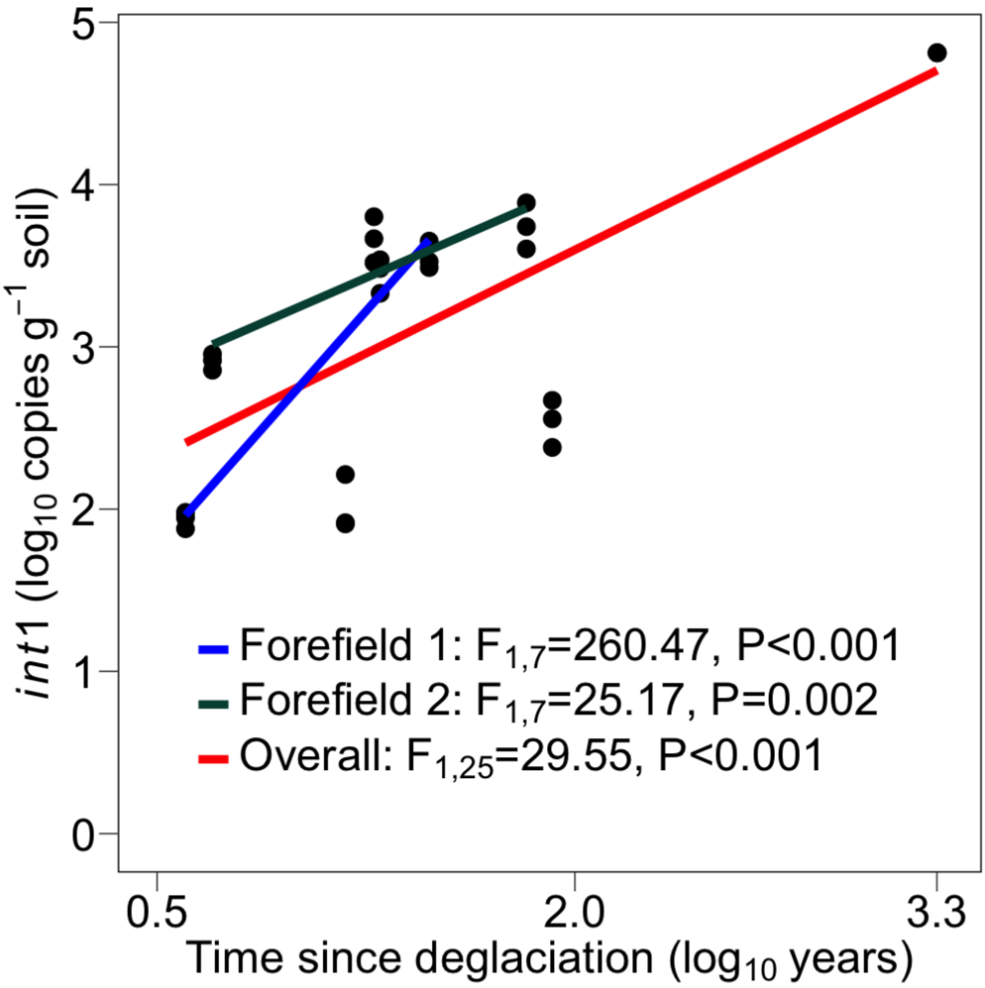


Figure S3. Relationship assessed as regression between chronosequence (time since deglaciation), represented as a continuous variable, and mobile genetic element (*int*1 abundance) for two glacier forefields and overall, all samples together (red line). Forefield 1 (blue line): Austre Brøggerbreen glacier forefield contains samples MHG1, MHG2, MHG3; Forefield 2 (green line): Midtre Lovénbreen glacier forefield contains samples MHG4, MHG5, MHG6.

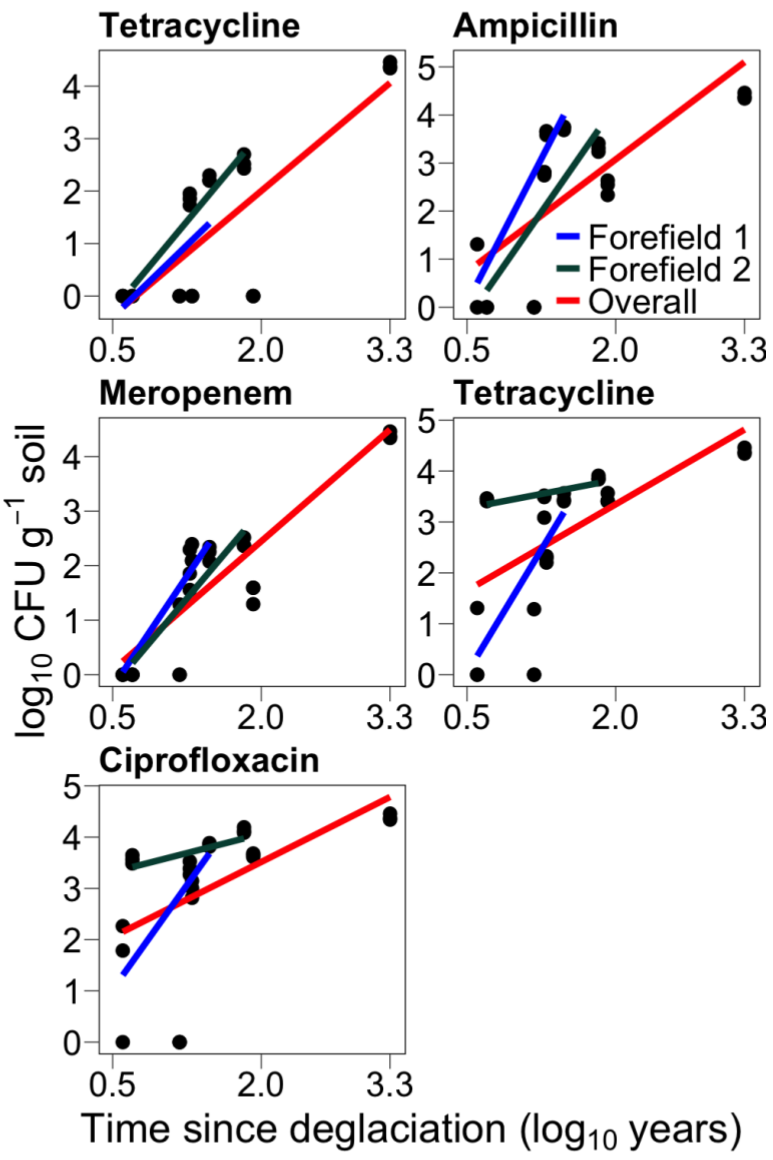


Figure S4. Relationship assessed as regression between chronosequence (time since deglaciation), represented as a continuous variable, and abundance of antibiotic resistant bacteria when exposed to five different antibiotics for two glacier forefields and overall, all samples together (red line). The associated statistics are described in Table S3. If the relationship is not significant for any of the combinations, then the regression line (coloured lines) is absent. Forefield 1 (blue line): Austre Brøggerbreen glacier forefield contains samples MHG1, MHG2, MHG3; Forefield 2 (green line): Midtre Lovénbreen glacier forefield contains samples MHG4, MHG5, MHG6.

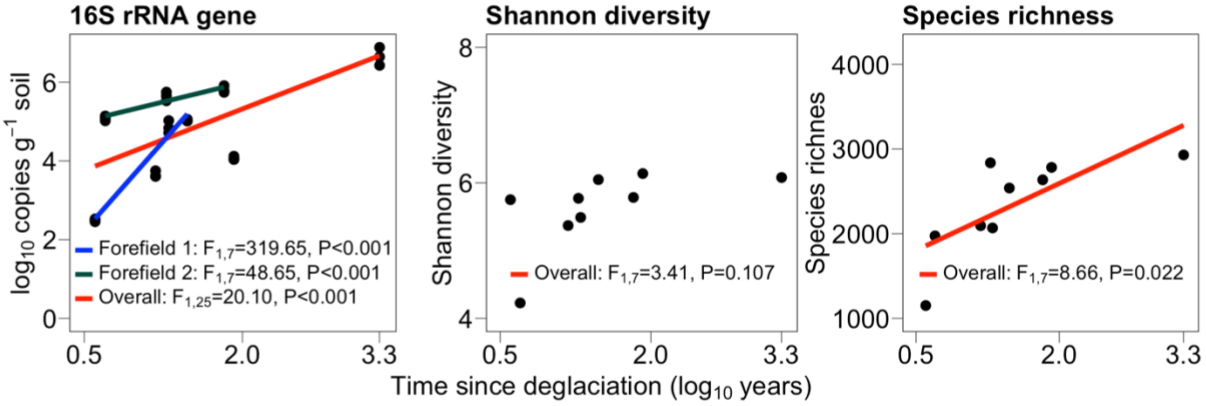


Figure S5. Relationship assessed as regression between chronosequence (time since deglaciation), represented as a continuous variable, and microbial diversity for two glacier forefields and overall, all samples together (red line). Microbial diversity is evaluated as 16S rRNA gene abundance, species richness and Shannon diversity. Separate linear models for different forefields were not evaluated for Shannon diversity and species richness. Forefield 1 (blue line): Austre Brøggerbreen glacier forefield contains samples MHG1, MHG2, MHG3; Forefield 2 (green line): Midtre Lovénbreen glacier forefield contains samples MHG4, MHG5, MHG6.

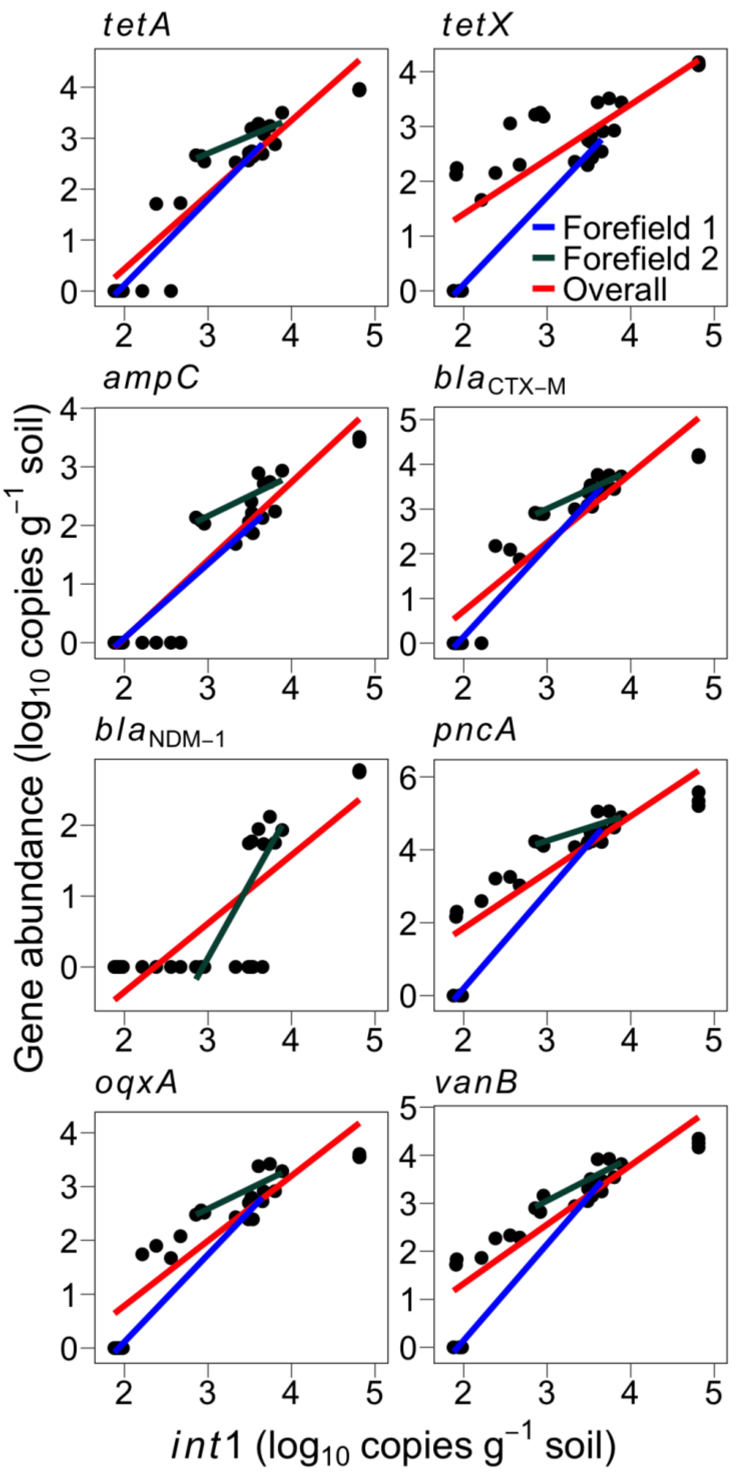


Figure S6. Relationship assessed as regression between the mobile genetic element (*int*1 abundance) and abundance of eight ARGs expressed as copies per gram soil for two glacier forefields and overall all samples together (red line). The associated statistics are described in Table S4. If the relationship is not significant for any of the combinations, then the regression line (coloured lines) is absent. Forefield 1 (blue line): Austre Brøggerbreen glacier forefield contains samples MHG1, MHG2, MHG3; Forefield 2 (green line): Midtre Lovénbreen glacier forefield contains samples MHG4, MHG5, MHG6.

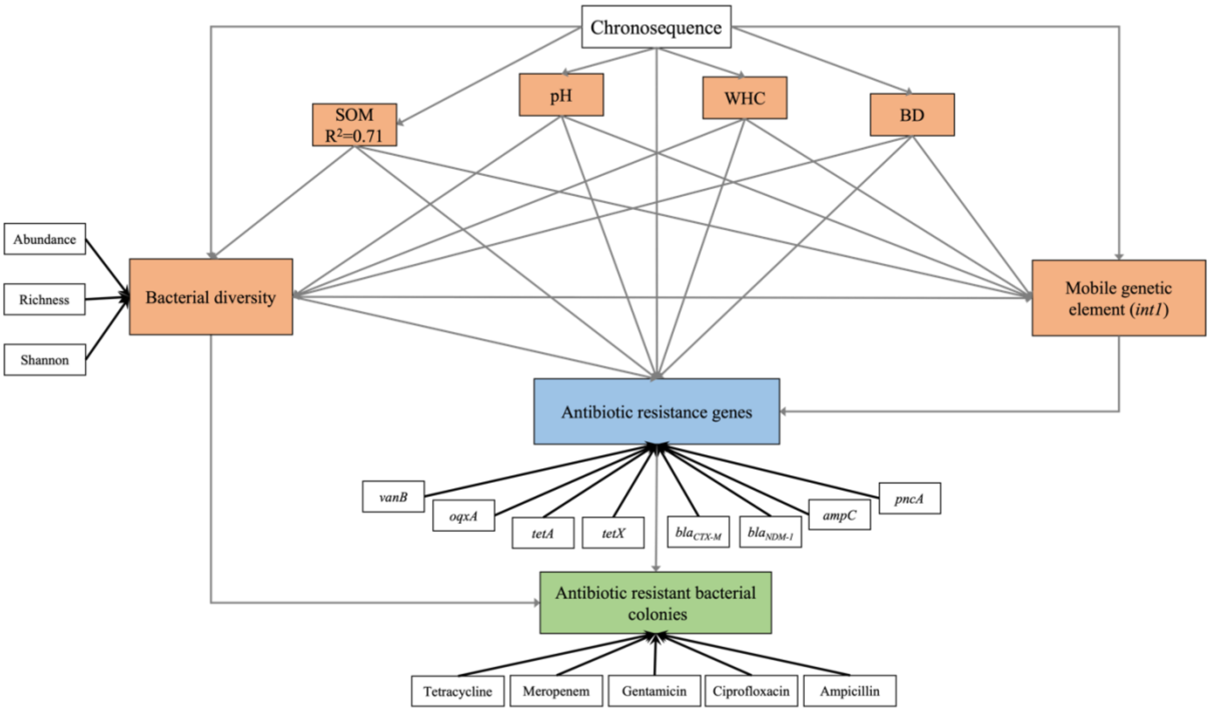


Figure S7. Partial least squares path model (PLS-PM) showing all modelled paths.

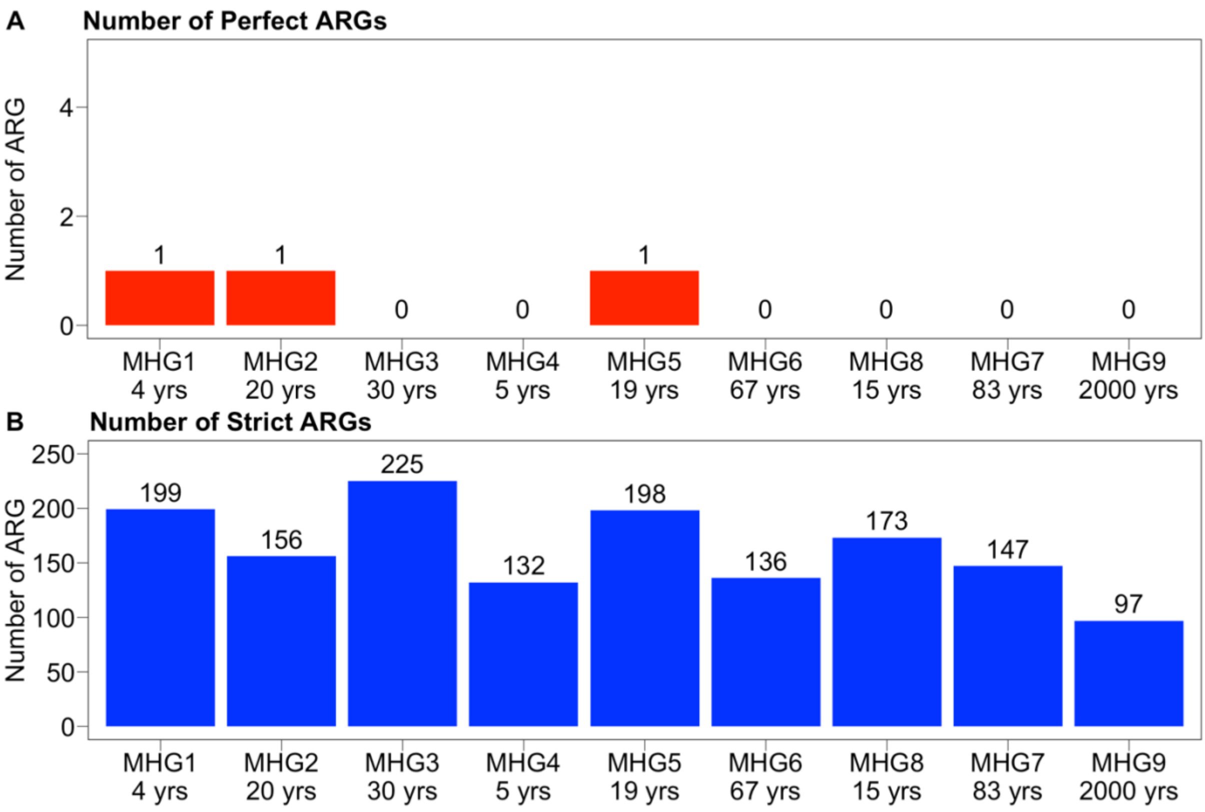


Figure S8. Number of ARGs identified from metagenomes further classified into Perfect hits (A) and Strict hits (B) along a chronosequence of deglaciated soils at different glacier forefields.

Table S1. Summary of ANOVA results evaluating the variation of individual ARGs (expressed as copies $g^{-1}$ soil) with soil age. See Fig S1.

| ARG | Overall | | Forefield 1 | | Forefield 2 | |
|---|---|---|---|---|---|---|
| | F-value | P-value | F-value | P-value | F-value | P-value |
| *tetA* | $F_{1,25}$=10.77 | 0.003 | $F_{1,7}$=317.27 | <0.001 | $F_{1,7}$=53.34 | <0.001 |
| *tetX* | $F_{1,25}$=20.18 | <0.001 | $F_{1,7}$=744.19 | <0.001 | $F_{1,7}$=1.16 | 0.317 |
| *ampC* | $F_{1,25}$= 9.23 | 0.006 | $F_{1,7}$=832.60 | <0.001 | $F_{1,7}$=43.97 | <0.001 |
| $bla_{CTX\text{-}M}$ | $F_{1,25}$=11.65 | 0.002 | $F_{1,7}$=604.23 | <0.001 | $F_{1,7}$=130.38 | <0.001 |
| $bla_{NDM\text{-}1}$ | $F_{1,25}$=28.48 | <0.001 | $F_{1,7}$=2.28 | 0.175 | $F_{1,7}$=20.20 | 0.003 |
| *pncA* | $F_{1,25}$=13.68 | 0.001 | $F_{1,7}$=210.01 | <0.001 | $F_{1,7}$=162.06 | <0.001 |
| *vanB* | $F_{1,25}$=16.98 | <0.001 | $F_{1,7}$=405.94 | <0.001 | $F_{1,7}$=108.73 | <0.001 |
| *oqxA* | $F_{1,25}$=13.69 | 0.001 | $F_{1,7}$=894.86 | <0.001 | $F_{1,7}$=227.75 | <0.001 |

Table S2. Summary of ANOVA results evaluating the variation of relative abundance of individual ARGs (expressed as copies per copy of 16S rRNA gene) with soil age. See Fig S2.

| ARG (relative abundance) | Overall | | Forefield 1 | | Forefield 2 | |
|---|---|---|---|---|---|---|
| | F-value | P-value | F-value | P-value | F-value | P-value |
| *tetX* | $F_{1,25}$=6.99 | 0.014 | $F_{1,7}$=96.25 | <0.001 | $F_{1,7}$=2.55 | 0.155 |
| *tetA* | $F_{1,25}$=5.41 | 0.028 | $F_{1,7}$=173.62 | <0.001 | $F_{1,7}$=0.01 | 0.921 |
| *ampC* | $F_{1,25}$=2.12 | 0.158 | $F_{1,7}$=106.76 | <0.001 | $F_{1,7}$=0.06 | 0.813 |
| $bla_{CTX\text{-}M}$ | $F_{1,25}$=9.93 | 0.004 | $F_{1,7}$=80.77 | <0.001 | $F_{1,7}$=3.94 | 0.088 |
| $bla_{NDM\text{-}1}$ | $F_{1,25}$=14.76 | <0.001 | $F_{1,7}$=2.28 | 0.175 | $F_{1,7}$=14.49 | 0.007 |
| *pncA* | $F_{1,25}$=7.76 | 0.010 | $F_{1,7}$=75.44 | <0.001 | $F_{1,7}$=0.37 | 0.563 |
| *vanB* | $F_{1,25}$=9.49 | 0.005 | $F_{1,7}$=97.68 | <0.001 | $F_{1,7}$=2.42 | 0.164 |
| *oqxA* | $F_{1,25}$=9.85 | 0.004 | $F_{1,7}$=109.12 | <0.001 | $F_{1,7}$=0.61 | 0.459 |

Table S3. Summary of ANOVA results evaluating the variation of ARB for individual antibiotics (expressed as CFU $g^{-1}$ soil) with soil age. See Fig S4.

| | Overall | | Forefield 1 | | Forefield 2 | |
|---|---|---|---|---|---|---|
| | F-value | P-value | F-value | P-value | F-value | P-value |
| Tetracycline | $F_{1,25}=41.77$ | <0.001 | $F_{1,7}=5.27$ | 0.055 | $F_{1,7}=108.57$ | <0.001 |
| Ampicillin | $F_{1,25}=27.34$ | <0.001 | $F_{1,7}=83.29$ | <0.001 | $F_{1,7}=49.74$ | <0.001 |
| Meropenem | $F_{1,25}=93.39$ | <0.001 | $F_{1,7}=162.56$ | <0.001 | $F_{1,7}=53.32$ | <0.001 |
| Gentamicin | $F_{1,25}=14.99$ | <0.001 | $F_{1,7}=52.66$ | <0.001 | $F_{1,7}=6.25$ | 0.041 |
| Ciprofloxacin | $F_{1,25}= 9.72$ | 0.005 | $F_{1,7}=21.11$ | 0.003 | $F_{1,7}=6.79$ | 0.035 |
| No antibiotic | $F_{1,25}=15.61$ | <0.001 | $F_{1,7}=850.61$ | <0.001 | $F_{1,7}=5.47$ | 0.052 |

Table S4. Summary of ANOVA results evaluating the variation of individual ARGs (expressed as copies $g^{-1}$ soil) with MGE (*int*1 abundance; copies $g^{-1}$ soil). See Fig S6.

| ARG | Overall | | Forefield 1 | | Forefield 2 | |
|---|---|---|---|---|---|---|
| | F-value | P-value | F-value | P-value | F-value | P-value |
| *tetX* | $F_{1,25}$=42.03 | >0.001 | $F_{1,7}$=348.66 | >0.001 | $F_{1,7}$=0.01 | 0.976 |
| *tetA* | $F_{1,25}$=148.30 | >0.001 | $F_{1,7}$=959.01 | >0.001 | $F_{1,7}$= 1.83 | 0.004 |
| *ampC* | $F_{1,25}$=165.78 | >0.001 | $F_{1,7}$=596.20 | >0.001 | $F_{1,7}$= 1.15 | 0.012 |
| $bla_{CTX-M}$ | $F_{1,25}$=133.71 | >0.001 | $F_{1,7}$=811.00 | >0.001 | $F_{1,7}$= 4.61 | >0.001 |
| $bla_{NDM-1}$ | $F_{1,25}$=43.53 | >0.001 | $F_{1,7}$=1.17 | 0.315 | $F_{1,7}$= 2.05 | 0.003 |
| *pncA* | $F_{1,25}$=74.17 | >0.001 | $F_{1,7}$=809.67 | >0.001 | $F_{1,7}$= 1.47 | 0.006 |
| *vanB* | $F_{1,25}$=96.05 | >0.001 | $F_{1,7}$=1044.28 | >0.001 | $F_{1,7}$= 2.67 | 0.001 |
| *oqxA* | $F_{1,25}$=109.3 | >0.001 | $F_{1,7}$=610.44 | >0.001 | $F_{1,7}$= 1.62 | 0.005 |

Table S5. A-priori evidence in the literature for the modelled paths in PLS-PM.

| **Path** | **References** |
|---|---|
| Chronosequence → SOM | (*16*, *17*) |
| Chronosequence → pH | (*18*) |
| Chronosequence → WHC | (*18*) |
| Chronosequence → BD | (*18*) |
| Chronosequence → Bacterial diversity | (*19*, *20*) |
| Chronosequence → Antibiotic resistance genes | (*21*) |
| Chronosequence → Mobile genetic element | (*21*) |
| SOM → Bacterial diversity | (*19*, *20*) |
| SOM → Antibiotic resistance genes | (*21*) |
| SOM → Mobile genetic element | (*21*) |
| pH → Bacterial diversity | (*22*) |
| pH → Antibiotic resistance genes | (*23*, *24*) |
| pH → Mobile genetic element | (*23*, *24*) |
| WHC → Bacterial diversity | (*25*) |
| WHC → Antibiotic resistance genes | (*24*) |
| WHC → Mobile genetic element | (*24*) |
| BD → Bacterial diversity | (*25*) |
| BD → Antibiotic resistance genes | (*24*) |
| BD → Mobile genetic element | (*24*) |
| Bacterial diversity → Antibiotic resistance genes | (*26*) |
| Bacterial diversity → Mobile genetic element | (*26*) |
| Mobile genetic element → Antibiotic resistance genes | (*27*) |
| Bacterial diversity → Antibiotic resistant bacterial colonies | (*26*) |
| Antibiotic resistance genes → Antibiotic resistant bacterial colonies | (*28*) |

Table S6. Soil edaphic factors from sample sites in deglaciated forefields.

| Sample | Time since deglaciation till sampling (years) | Distance from glacier snout (m) | Soil organic matter (SOM; %) | pH | Bulk density (BD; g $cm^{-3}$) | Water holding capacity (WHC; %) |
|---|---|---|---|---|---|---|
| MHG1 | 4 | 76 | 0.401 ± 0.726 | 6.48 | 1.32 | 47.47 |
| MHG2 | 20 | 388 | 0.729 ± 0.231 | 7.51 | 1.28 | 40.65 |
| MHG3 | 30 | 594 | 1.577 ± 0.685 | 6.1 | 1.28 | 37.80 |
| MHG4 | 5 | 43 | 1.014 ± 0.467 | 5.74 | 1.03 | 52.59 |
| MHG5 | 19 | 137 | 1.857 ± 1.023 | 5.25 | 1.23 | 54.04 |
| MHG6 | 67 | 850 | 2.560 ± 0.330 | 5.34 | 1.24 | 44.44 |
| MHG7 | 83 | 2000 m (from Baronbreen) | 0.807 ± 0.039 | 8 | 1.23 | 27.96 |
| MHG8 | 15 | 180 | 2.635 ± 0.172 | 5.07 | 1.19 | 42.18 |
| MHG9 | 2000 | NA | 48.135 ± 2.613 | 6.3 | 1.53 | 52.21 |

Table S7. Summary of antibiotic classes chosen in the study

| Antibiotic class | Common antibiotics | Mechanism of action |
|---|---|---|
| Tetracycline | **Tetracycline**, Doxycycline, Minocycline | Broad-spectrum bacteriostatic antibiotics that bind to the 30S ribosomal subunit and prevent the aminoacyl tRNA binding to the A site of the ribosome. |
| Beta-lactam | Penicillin, **Ampicillin**, Amoxicillin, Cephalosporin, **Meropenem** (Carbapenems) | Bactericidal antibiotics that inhibit the synthesis of the peptidoglycan layer of bacterial cell walls. The antibiotic covalently binds to the essential penicillin-binding proteins (PBPs), which are enzymes involved in the bacterial peptidoglycan transpeptidation. |
| Aminoglycoside | **Gentamicin**, Streptomycin, Amikacin | Narrow-spectrum bactericidal antibiotics that inhibit peptide elongation at the 30S ribosomal subunit during protein synthesis. |
| Pyrazinamide | Pyrazinamide | Pyrazinamide is a widely used prodrug that prevents the growth of *Mycobacterium tuberculosis*. |
| Quinolone | **Ciprofloxacin**, Ofloxacin | Broad-spectrum bactericidal antibiotics that inhibit the ligase activity of the type II topoisomerases, DNA gyrase and topoisomerase IV, thereby preventing DNA replication. |
| Glycopeptide | Vancomycin | Multiple mechanisms. Vancomycin prevents the addition of new units to the peptidoglycan of bacterial cell walls. |

Table S8. Primers for 16S rRNA gene and antibiotic-resistant genes (ARGs).

| Target genes | Primer | Sequences (5′—3′) | Antibiotic class | Reference |
|---|---|---|---|---|
| *16S rRNA* | *F515* | GTGCCAGCMGCCGCGGTAA | | (*9*) |
| | *R806* | GGACTACVSGGGTATCTAAT | | |
| *tetX* | *tet*X-F | CAATAATTGGTGGTGGACCC | Tetracycline | (*10*) |
| | *tet*X-R | TTCTTACCTTGGACATCCCG | | |
| *tetB/P* | *tet*B/P-F | AAAACTTATTATATTATAGTG | Tetracycline | (*11*) |
| | *tet*B/P-R | TGGAGTATCAATAATATTCAC | | |
| *tetA* | *tetA*-F | CTCACCAGCCTGACCTCGAT | Tetracycline | (*11*) |
| | *tetA*-R | CACGTTGTTATAGAAGCCGCATAG | | |
| *ampC* | *amp*-F | GCAGCACGCCCCGTAA | Ampicillin (β-lactam) | (*11*) |
| | *amp*-R | TGTACCCATGATGCGCGTACT | | |
| $bla_{CTX-M}$ | *CTX*-F | GGAGGCGTGACGGCTTTT | β-lactam | (*11*) |
| | *CTX*-R | TTCAGTGCGATCCAGACGAA | | |
| $bla_{IMP}$ | *IMP*-F | AACACGGTTTGGTGGTTCTTGTA | β-lactam | (*11*) |
| | *IMP*-R | GCGCTCCACAAACCAATTG | | |
| $bla_{NDM-1}$ | *NDM*-F | ATTAGCCGCTGCATTGAT | β-lactam | (*12*) |
| | *NDM*-R | CATGTCGAGATAGGAAGTG | | |
| $bla_{OXA}$ | *OXA*-F | CGGATGGTTTGAAGGGTTTATTAT | β-lactam | (*11*) |
| | *OXA*-R | TCTTGGCTTTTATGCTTGATGTTAA | | |
| $bla_{TEM}$ | *TEM*-F | AGCATCTTACGGATGGCATGA | β-lactam | (*11*) |
| | *TEM*-R | TCCTCCGATCGTTGTCAGAAGT | | |
| *AAC(3)-IIc or aacC2* | *aacC2*-F | ACGGCATTCTCGATTGCTTT | Aminoglycoside | (*11*) |

| | | | | |
|---|---|---|---|---|
| | *aacC2R* | CCGAGCTTCACGTAAGCATTT | | |
| *pncA* | *pncA*-F | GCAATCGAGGCGGTGTTC | Pyrazinamide | (*13*) |
| | *pncA*-R | TTGCCGCAGCCAATTCA | | |
| *intI1* | *intI1*-F | CGAACGAGTGGCGGAGGGTG | Integron | (*14*) |
| | *intI1*-R | TACCCGAGAGCTTGGCACCCA | | |
| *tnpA*-01/*tnpA* | *tnpA*-F | CATCATCGGACGGACAGAATT | Transposase-IS21 Group | (*14*) |
| | *tnpA*-R | GTCGGAGATGTGGGTGTAGAAAGT | | |
| *vanA* | *vanA*-F | AAAAGGCTCTGAAAACGCAGTTAT | Vancomycin (Glycopeptide) | (*11*) |
| | *vanA*-R | CGGCCGTTATCTTGTAAAAACAT | | |
| *vanB* | *vanB*-F | TTGTCGGCGAAGTGGATCA | Vancomycin (Glycopeptide) | (*11*) |
| | *vanB*-R | AGCCTTTTTCCGGCTCGTT | | |
| *oqxA* | *oqxA*-F | GACAGCGTCGCACAGAATG | Quinolone, detergent, trimethoprim | (*15*) |
| | *oqxA*-R | GGAGACGAGGTTGGTATGGA | | |
